# Controllable Fusion of Human Brain Organoids Using Acoustofluidics

**DOI:** 10.1101/2020.08.05.238113

**Authors:** Zheng Ao, Hongwei Cai, Zhuhao Wu, Jonathan Ott, Huiliang Wang, Ken Mackie, Feng Guo

## Abstract

The fusion of human organoids holds promising potential in modeling physiological and pathological processes of tissue genesis and organogenesis. However, current fused organoid models face challenges of high heterogeneity and variable reproducibility, which may stem from the random fusion of heterogeneous organoids. Thus, we developed a simple and versatile acoustofluidic method to improve the standardization of fused organoid models via a controllable spatial arrangement of organoids. By regulating dynamic acoustic fields within a hexagonal acoustofluidic device, we can rotate, transport, and fuse one organoid with another in a contact-free, label-free, and minimal-impact manner. As a proof-of-concept to model ventral tegmentum (VTA)-prefrontal cortex (PFC) projection, we acoustically fused human forebrain organoids (hFOs) and human midbrain organoids (hMOs) with the controllable alignment of neuroepithelial buds. We characterized the successful development of fused assembloids via robust tyrosine hydroxylase (TH) neuron projection, accompanied by an increase of firing rates and synchrony of excitatory neurons. Moreover, we found that our controllable fusion can promote neuron projection (e.g., range, length, and density), projection maturation (e.g., higher firing rate and synchrony), and neural progenitor cell (NPC) division in the assembloids. Thus, our acoustofluidic method would facilitate the standardization and robustness of organoid-based disease models and tissue engineering.

---

Human organoids, or organ-like 3D cultures, are formed through the spontaneous self-organization and differentiation of human pluripotent stem cells (hPSCs) or human tissue-derived progenitor cells.[1] Human organoid cultures provide a promising platform to recapitulate key histological features of human healthy and diseased organs, holding great potential in organ developmental research, disease modeling, drug screening, and translational medicine. Compared with 2D in vitro cell cultures, the human organoid cultures recapitulate key physiological features and complexity of human tissues and organs. Compared with animal models, human organoids represent human-specific genetics, epigenetics, and transcriptomics. So far, a variety of human organoids have been developed for optic cup, intestine, kidney, lung, pancreas, thymus, various types of tumors as well as brain tissues.[2] Moreover, the fused human organoids have been explored to recapitulate the complete organ makeup or to study inter-organ cross-talks.[3] For example, brain assembloids, the cultures of fused human brain region-specific organoids, have been generated to model complex cell-cell interactions and neural circuit formation (e.g., cortico-striatal, cortical-spinal, meso-cortical, and cortico-hippocampal circuits) in the human nervous system.[4–11] Additionally, brain organoids can also be fused with glioblastoma spheroids to model tumorigenesis and invasion.[12] The fused human organoid models have tremendous potential in etiology and drug development with multi-organ crosstalk modeling including brain-colorectal assembloids to model brain-gut axis, brain-pituitary gland organoids to model neuro-endocrine axis, etc. However, high heterogeneity and variable reproducibility limit the application potentials of current fused human organoid models. Current human organoids have high heterogeneity in size, shape, and spatial distribution of asymmetric features such as neuroepithelial buds in brain organoids. The random fusion of organoids without controlling the distribution and position of these asymmetric features may result in highly diverse inter-organoid cell migration/projection dynamics and neurological function.

Recently, efforts have been made to address the technological challenges in generating robust, reproducible, and reliable human organoid and fused human organoid models. One strategy is to generate standardized human organoids as well as improve the quality control of organoid cultures. For example, Paşca group developed a directed differentiation method for the reliable generation of human cortical organoids derived from human induced pluripotent stem cells (hiPSCs) by modulating the SMAD and Wnt pathways and the growth factors EGF and FGF2.[13] Lutolf and his colleagues employed microengineered devices and automated systems for high throughput generation of gastrointestinal organoids with reduced heterogeneity.[14] Qin research team took advantage of droplet microfluidics to achieve the reproducible and high-throughput fabrication of uniform islet organoids derived from hiPSCs.[15] Additional to improve the uniformity or standardization of human organoids, a controllable fusion of asymmetric human organoids could be an attractive alternative solution to further reduce the heterogeneity of fused human organoid and assembloid models. Recently, our group developed an acoustofluidics technology that employs microfluidics and acoustics for the generation and manipulation of 3D cell spheroids and organoids. Superior to other technologies that employ magnetic, dielectronic, and/or optical forces, this acoustofluidics technology can achieve the rational and transportational manipulation of biological specimens from micro to macro-scale in a label-free, contact-free, and highly biocompatible manner, preserving cell cultures in their native medium.[16–18] Thus, our acoustofluidics technology may provide a unique solution for simple, versatile, and effective engineering to reproducibly generate fused human organoids for various applications through the controllable fusion of human organoids with heterogeneous morphological and structural features.

Here, using our acoustofluidic technique, we aim to study the dopaminergic pathway in an in vitro human brain environment through the controllable fusion of human forebrain organoids (hFOs) and human midbrain organoids (hMOs) into hFO-hMO assembloids. Dopaminergic pathways regulate learning and reward-related behaviors, and the dysfunction of this pathway during early brain development can lead to neurological diseases such as schizophrenia, attention-deficit hyperactivity disorder (ADHD), and autism spectrum disorder (ASD).[19] The mesocortical pathway includes dopaminergic projections from the midbrain ventral tegmentum (VTA) to the prefrontal cortex (PFC).[20] It is of great interest to establish an *in vitro* human model of dopaminergic projections from VTA to PFC to reliably model mesocortical pathway formation during early neural development.[21] In a developing human brain, dopamine neurons project from VTA toward PFC during early development (**Figure 1a**). To mimic the VTA-PFC projection, we generated hFOs and hMOs (**Figure S1-2**) derived from human embryonic stem cells (hESC, WA01) using an adopted brain region-specific organoids fabrication protocol (**Table S1-2**).[22] These hFOs and hMOs showed vast heterogeneity in size, morphology, distribution of asymmetric features such as neuroepithelial buds (referred to as “bud” in following denotations). For example, we observed 18-day-old hFOs with highly variable sizes and shapes (size ranging from 11,543 μm_2_ to 54,508 μm_2_; roundness ranging from 0.37 to 0.98; n=40, **Figure S3a**). Moreover, we also found the number, size, location, and orientation of neuroepithelial buds (indicated by yellow circles) vary dramatically in-between organoids. This indicated the uneven distribution of underlying VZ/SVZ regions within these organoids as they align with bud distribution (**Figure 1b**). We analyzed these hFOs to quantify the variation of bud sizes and orientation (orientation angle α, 119.0 ± 101.2o; bud size, 4654 ± 3617 μm_2_; n=40, **Figure S3b**). Given such heterogeneity, the uncontrolled fusion of heterogeneous hFOs and hMOs will likely introduce significant heterogeneity to the formation of hFO-hMO assembloids.

**Figure 1.**
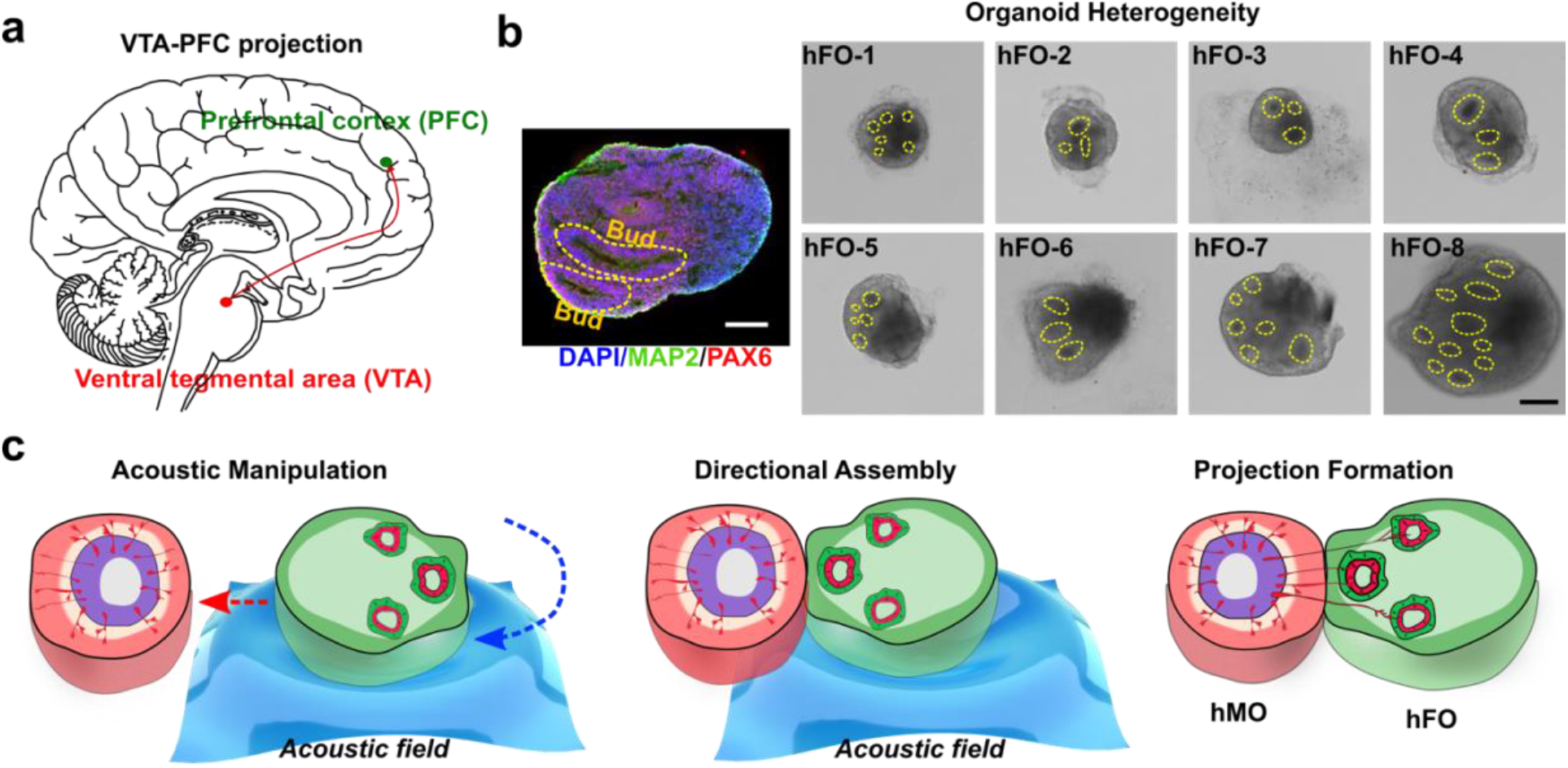
The controllable fusion of hMOs and hFOs to model VTA-PFC projections. (a) Representation of VTA-PFC projection in a human brain. (b) Heterogeneity of human forebrain organoids (hFOs). Eight hFOs generated from one bench have the random number and location of neuroepithelial buds (in yellow circle). The neuroepithelial bud presents the structure of the subventricular zone (neural progenitor marker PAX6, in red; neuron marker MAP2, green; cell nucleus DAPI, blue). (c) The acoustic fusion of a hMO and a hFO into a hMO-hFO assembloid with precision control over organoid arrangement and neuroepithelial bud location. Scale bar: 100 μm

Thus, we developed an acoustofluidic method to manipulate and fuse organoids with excellent control over the spatial arrangement of hFO and hMO for better regulation of neuron projection. By regulating dynamic acoustic fields within a hexagonal acoustofluidic device, we can rotate, transport, and fuse one hFO with one hMO in a contact-free, label-free, and highly-biocompatible manner (**Figure 1c**). We developed the acoustofluidic method for the organoid manipulation by incorporating dynamically-regulated acoustic fields within a hexagonal acoustofluidic device. The hexagonal acoustofluidic device consists of pairs of piezoelectric transducers (PZT, lead zirconate titanate), and an organoid chamber (**Figure 2a**). By independently tuning the phase and amplitude of input radio frequency (RF) signals, the generated dynamic pattern of acoustic fields can enable the rotational and transportational manipulation of single organoids. To predict and optimize human organoid handling, we simulated the acoustic fields generated within the hexagonal acoustofluidic device. By sequentially increasing the amplitude of RF signals applied to one selected PZT pair within the three pairs, the orientation of periodically-distributed pressure nodes (blue nodes) rotated accordingly in a clockwise direction (**Figure 2a**). By tuning the phase-angle between two PZT transducers with the selected pair, the position of periodically-distributed pressure nodes (blue nodes) moved along the PZT pair direction forward or backward accordingly (**Figure 2a**). Then, after trapping a hFO within pressure nodes in our hexagonal acoustofluidic device, we rotated the trapped organoid from 0o to 60 o, 120 o, 180 o, 240 o, and 300 o (**Figure 2b**) by sequentially rotating the orientation of pressure node patterns. We further quantified the rotational speed of the trapped organoid by modulating the amplitudes of RF signals applied to the three PZT pairs (Pair1 = 6.8 Vpp, Pair2 = 6.8 Vpp, Pair3 = 10 ~ 40 Vpp). By increasing the amplitude of PZT pair, the rotational speed of the trapped organoid increased accordingly (**Figure 2c**). Moreover, our method can achieve the on-demand rotation angle by controlling the time and amplitudes of the applied RF signals to the three PZT pairs. Similarly, by modulating the amplitude, phase-angle, and duration of RF signals applied to the three PZT pairs, the trapped organoid can be transported to the desired distance along the PZT pair direction with a moving speed ranging from 0 to 3 mm/s (**Figure 2d**). Thus, by combining the rational and transportational manipulation of organoids, we can trap, move, and/or rotate the selected organoid to target location with desired orientation within our organoid chamber.

**Figure 2.**
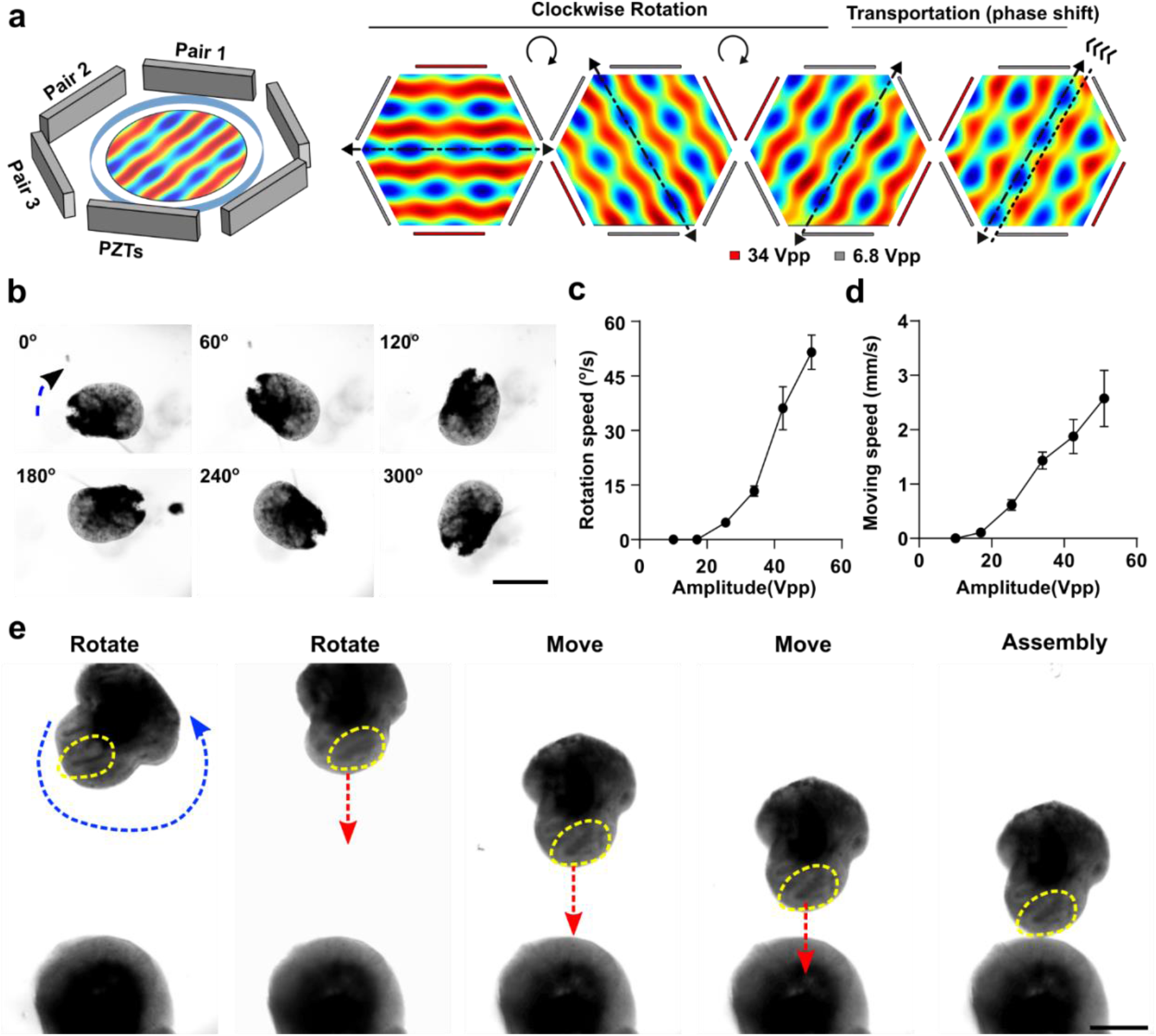
Acoustic manipulation and assembly of organoids. (a) Schematics of the acoustic device, and simulation of Gor’kov potential fields for the rotational and transportational manipulation of organoids. (b) Corresponding experimental results (optical microscopy images) taken for the three amplitudes combinations. The organoid was trapped along 30, 90, 120, 180, 240, and 300 degrees (to the vertical direction) respectively. (c) Experimental results showing the dependence of the rotational angular speeds on the input amplitudes. (d) Experimental results showing the dependence of the manipulation movement speed on the input amplitudes. (e) Demonstration of acoustic directional assembly of hFO and hMO. Scale bars, 500 μm

We further applied our acoustofluidic method for the directional fusion of hFOs and hMOs. By controlling the initial spatial arrangement of hFOs and hMOs, we expect to manipulate the arrangement and location of neuroepithelial buds in fused organoids, for example, to minimize the initial distance between hFO neuroepithelial buds and hMO. We first rotated the hFO via amplitude modulation, the neuroepithelial bud (indicated by yellow dotted circles) within the hFO was rotated to face toward to the hMO. Then, the organoids were transported along vertical direction step by step towards the immobilized hMO in the organoid chamber while maintaining the same relative orientation of the neuroepithelial bud. By further tuning the phase, the hFO finally contacted with the hMO with the desired orientation for neuroepithelial bud alignment (**Figure 2e**). Furthermore, using this method, we could fine-tune the initial distance between the hFO neuroepithelial bud and hMO by precision control of hFO bud orientation. We first defined the distance between the hMO-hFO contact point and the geometric center of the hFO “bud” and as “d”, and its distance from the geometric center of the whole hFO as “r”. We manipulated the fusion of hFOs and hMOs to generate “*Toward* group” (d < r, short d) and “*Away* group” (d > r, long d, **Figure 3a**). We also confirmed our acoustic distance manipulation by comparing the median of d in both groups (*Toward* group, 121.3 μm; *Away* group, 391.5 μm; p < 0.001, **Figure 3b**). To note, during the fusion of hMOs and hFOs, we rotated the hFOs to ensure all buds on the same hFO face toward (or away) from the hMO in the *Toward* (or *Away*) group (**Figure S4**). In a few cases where hFO buds were more evenly distributed and could not be rotated into one side of the midline, we excluded those hFOs from the following analysis.

**Figure 3.**
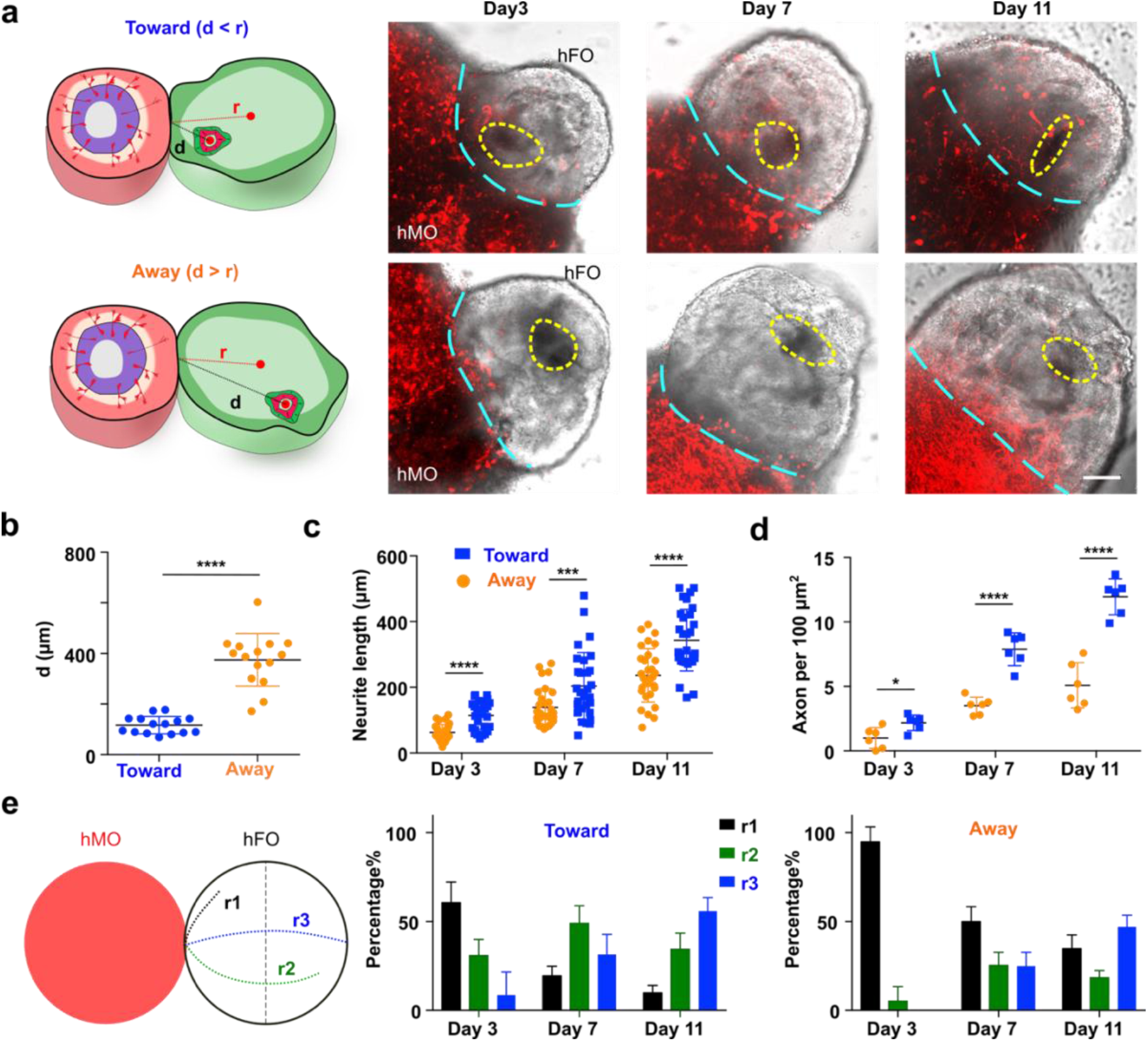
Fusion dynamics of hMO-hFO assembloids. (a) hFO was rotated to have all bud toward (d<r) or away (d>r) from hMO. hMO neuron projections labeled with mCherry expression (red) were imaged and recorded under a fluorescent microscope, buds are marked with dotted yellow circles, and the interface between hMO and hFO is marked with cyan dotted lines. (b) Comparison of hFO bud distance from hMO in the toward and away group. (n=15, p<0.001). (c) Distribution of projection neurite length over time (n=30, p<0.001) (d) Average axon density and its dynamic change in the toward and away groups over time (n=6, p<0.001). (e) Definition of range index in hMO-hFO projection. hMO neurite projections that did not reach midline of hFO are scored as r1; hMO neurite projections past midline of FO but did not reach the very end of hFO are scored as r2; hMO neurite projections that reached the very end of hFO are scored as r3. Distribution of range index and the dynamic change over time was different in the toward and away group (n=6). Scale bar: 100 μm

We then characterized the neuron projection dynamics right after the acoustic fusion of hMOs and hFOs with the controlled initial distance “d”. During the neuron projection formation processes, we characterized the projection neurite length, density, and range index in both the *Toward* and *Away* groups. We transfected hMOs with mCherry to track their projecting axons towards hFOs in the acoustically-fused organoids, and carefully compared the hMO-hFO projections in the *Toward* and *Away* groups (**Figure 3a**). We found that the *Toward* group has significantly longer neurites than the *Away* group (e.g., *Toward* group, 343.3 ± 93.8 μm; *Away* group, 236.5 ± 81.3 μm; p < 0.001, n = 30, at day 11, **Figure 3c**). In addition, the *Toward* group also has significantly higher neurite density than the *Away* group (e.g., *Toward* group, 11.9 ± 1.4 per 100 μm_2_; *Away* group, 5.1 ± 1.8 per 100 μm_2_; p < 0.001, n = 6, at day 11, **Figure 3d**). Furthermore, we quantified the range index[23] as following: projecting midbrain neurons that did not cross the midline of hFOs were scored with a range index of r1; neurons crossing midline but did not reach the furthest end of hFOs were scored as r2; neurons projected to the very end of hFOs were scored as r3. We observed that within the *Toward* group, 31.5 ± 11.3% neurons projected to the very end of hFOs (r3) at day 7, and 55.8% ± 7.7% neurons scored r3 at day 11. In contrast, in the *Away* group only 25.0% ± 7.1% reached r3 by day 7 and 47.0% ± 6.6% at day 11 (**Figure 3e**). To conclude, we observed that the *Toward* group has the greater length, higher density, and faster formation of projection neurites than the *Away* group, indicating our acoustic organoid fusion can manipulate the formation processes of neuron projections. This is likely due to the initial proximity as well as higher concentrations of TH neuron axonal guidance molecules.[24]

We further characterized the establishment of hMO-hFO projection and the controllability of our acoustofluidic method over its development. The tyrosine hydroxylase (TH) neurons are a key subtype of midbrain dopamine (mDA) neurons, which is born in the VTA and then projecting to the striatum and prefrontal cortex in a developing human brain.[25] Additionally, it was reported that TH neuron projections outside VZ/SVZ promote the proliferation of neural progenitor cells inside VZ/SVZ both prenatally and postnatally[26, 27] Thus, we validated the establishment of VTA-PFC projection in our hFO-hMO assembloids by staining the projection of TH neurons. We observed that TH neurons in our hFO-hMO assembloids could preferably project from hMO to the thin layer outside of VZ/SVZ of hFOs (**Figure 4a**), recapitulating the pattern seen *in vivo* in rodents and human.[28, 29] Meanwhile, we stained and analyzed the makeup of proliferative neural progenitor cells (EdU+) in hFOs, and hFO-hMO assembloids (hFO alone group, 36.2 ± 6.4%; *Toward* group, 79.8 ± 7.4%; *Away* group 50.0 ± 12.8%; p < 0.005, n = 6, **Figure 4b**). A significant increase of proliferative neural progenitor cells in the hFO-hMO assembloids versus hFOs alone, indicating the functional recapitulation of VTA-PFC projection. For the hFO-hMO assembloids, a significant increase of proliferative neural progenitor cells, as well as the TH neurons, was observed in the *Toward* group versus the *Away* group, indicating our acoustofluidic fusion can manipulate the development of the VTA-PFC projections.

**Figure 4.**
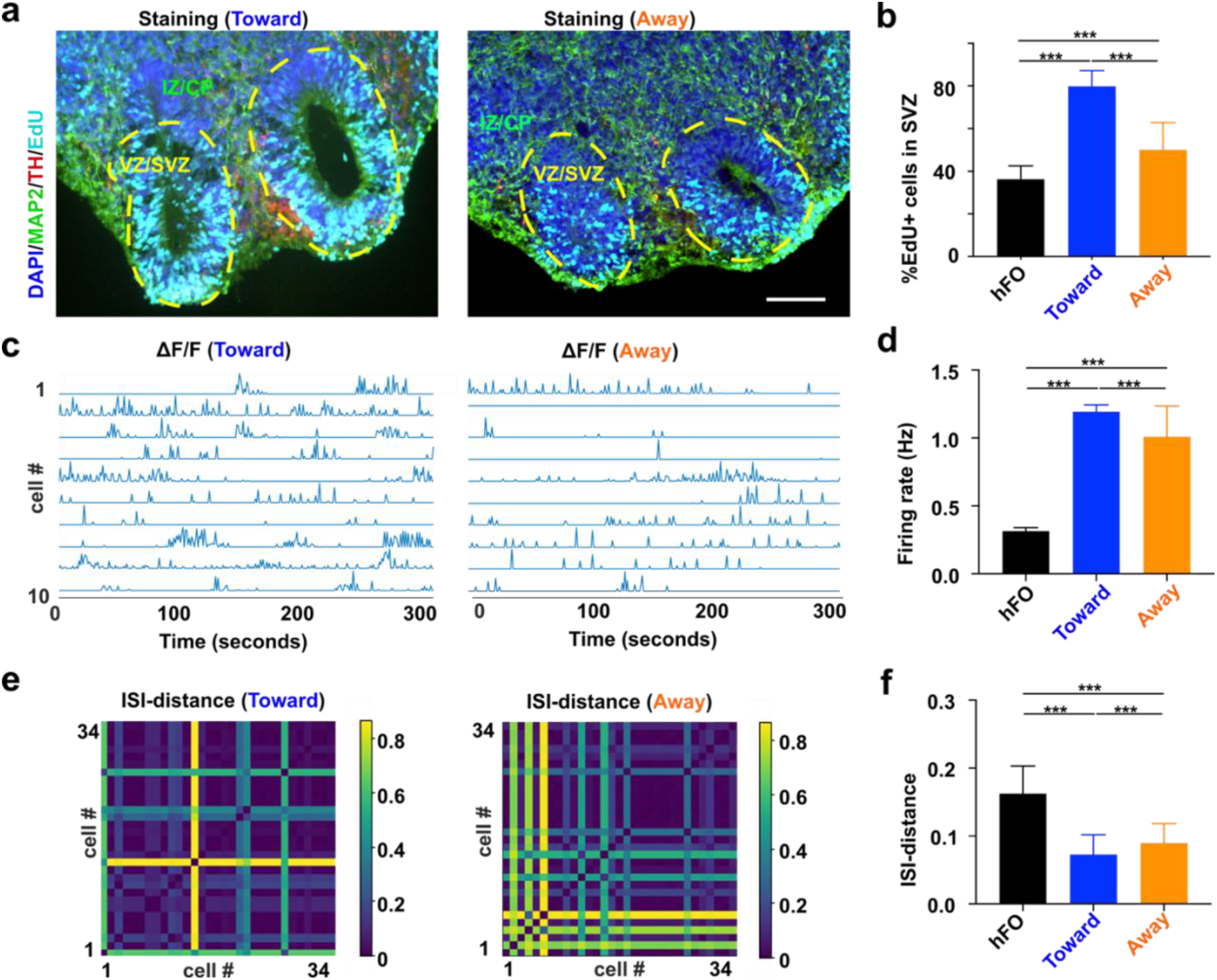
Control of projection formation in hMO-hFO assembloids. (a) Representative immunofluorescence image of hMO-hFO assembloid in the toward group (Cell nucleus-DAPI-blue, Dopamine neurons-TH-red, Forebrain neurons-MAP2-green, proliferative cells-EdU-cyan). (b) Percentage of newly proliferated cells (EdU+) cells in hFO, hMO-hFO assembloid in the toward and away group (p<0.005, n=6). (c) Representative calcium traces of ten active neurons extracted from movies of toward and away group. (d) Average firing rate of excitatory neurons in the hFO and assembloids in the toward and away groups. (e) ISI-distance matrix of 34 extracted calcium traces of excitatory neurons in the hFO and assembloids in the toward and away groups. (f) Average ISI-distance of excitatory neurons in hFO and the toward and away groups of hMO-hFO assembloids. Scale bar: 100 μm

Additionally, we also examined the electrophysiology of acoustically-fused hMO-hFO assembloids. *In vivo*, dopaminergic inputs from VTA was shown to modulate firing patterns and local field potential (LFP) oscillatory activity in PFC.[30] Activation of dopamine receptor-D1 positive PFC neurons also activates its burst firing, as indicated by higher firing rate and lower inter-spike intervals (ISIs).[31] To examine whether our hMO-hFO could recapitulate key in vivo electrophysiological features, we recorded and analyzed the spontaneous firing in hFOs and acoustically-fused hFO-hMO assembloids by calcium imaging. Briefly, hMO-hFO assembloids were transfected with adenovirus expressing GCaMP6s calcium reporter under a CAMKII promoter, limiting the calcium reporter expression in excitatory neurons in hFOs. We analyzed the spontaneous firing rate (**Figure 4c**) and ISI distances (**Figure 4e)** of excitatory neurons in 1-month-old hFO-hMO assembloids (*Toward* and *Away* groups), and hFOs (5 per group). They had significantly different spontaneous firing (hFOs alone, 0.31 ± 0.03 Hz, # of neurons = 63; *Toward* group, 1.19 ± 0.05 Hz, # of neurons = 312; *Away* group, 1.01 ± 0.23 Hz, # of neurons = 287; p < 0.005, **Figure 4d**) and ISI distances (hFOs alone, 0.16 ± 0.04, # of neurons = 63; *Toward* group, 0.073 ± 0.029, # of neurons = 312; *Away* group, 0.089 ± 0.029, # of neurons = 287; p < 0.005, **Figure 4f**). We found that (1) the hMO-hFO assembloids have significantly higher firing rate and synchrony (or lower ISI distance) than hFOs alone, indicating the formation of neuron projections; and (2) the *Toward* group of hMO-hFO assembloids have significantly higher firing rate and synchrony (or lower ISI distance) than the *Away* group, indicating that our acoustic organoid fusion method could manipulate the electrophysiology of fused organoids. Thus, we demonstrated we can mimic the key electrophysiological features of neuron projection development in the hMO-hFO assembloids, and our directional organoid fusion method can promote the maturation of neuronal network in hMO-hFO assembloids.

In summary, we have developed an acoustofluidic method to control the fusion of organoids for engineering robust and reliable fused human organoid and assembloid models. We demonstrated that the rotation and directional assembly of hFOs and hMOs could facilitate and manipulate the neuron projection and function in the hFO-hMO assembloids for modeling VTA-PFC projections. Taking the unique advantage of acoustics and microfluidics, this technique allows high-precision organoids or 3D in vitro cultures in a contact-free, label-free, highly compatible fashion. Moreover, this acoustofluidic technology is simple, versatile, and effective for engineering the fusion of organoids, which could be applicable for various organoids and 3D cultures with variable morphological features. We believe this tool can be broadly applied for human organoid-based organ developmental research, disease modeling, drug screening, and translational medicine.

## Experimental Section

See Supporting Information.

## Acknowledgment

Z.A. and H.C. contributed equally to this work. This project was supported by the departmental start-up funds of Indiana University Bloomington, and the National Institute of Health (1R03EB030331-01). H.W. acknowledges funding support from the National Institute of Health (1K01MH117490-01). The authors thank the Indiana University Imaging Center (NIH1S10OD024988-01) and Nanoscale Characterization Facility for use of their instruments.

## Conflict of Interest

The authors declare no conflict of interest.

## Notes

### Competing Interest Statement

The authors have declared no competing interest.

## References

1. Lancaster, M.A. and J.A. Knoblich, Organogenesis in a dish: Modeling development and disease using organoid technologies. Science, 2014. 345(6194): p. 1247125.

2. Schutgens, F. and H. Clevers, Human Organoids: Tools for Understanding Biology and Treating Diseases. Annu Rev Pathol, 2020. 15: p. 211–234.

3. Paşca, S.P., Assembling human brain organoids. Science, 2019. 363(6423): p. 126–127.

4. Pasca, S.P., The rise of three-dimensional human brain cultures. Nature, 2018. 553(7689): p. 437–445.

5. Jo, J., Y. Xiao, A.X. Sun, E. Cukuroglu, H.D. Tran, J. Goke, Z.Y. Tan, T.Y. Saw, C.P. Tan, H. Lokman, Y. Lee, D. Kim, H.S. Ko, S.O. Kim, J.H. Park, N.J. Cho, T.M. Hyde, J.E. Kleinman, J.H. Shin, D.R. Weinberger, E.K. Tan, H.S. Je, and H.H. Ng, Midbrain-like Organoids from Human Pluripotent Stem Cells Contain Functional Dopaminergic and Neuromelanin-Producing Neurons. Cell Stem Cell, 2016. 19(2): p. 248–257.

6. Qian, X., Ha N. Nguyen, Mingxi M. Song, C. Hadiono, Sarah C. Ogden, C. Hammack, B. Yao, Gregory R. Hamersky, F. Jacob, C. Zhong, K.-j. Yoon, W. Jeang, L. Lin, Y. Li, J. Thakor, Daniel A. Berg, C. Zhang, E. Kang, M. Chickering, D. Nauen, C.-Y. Ho, Z. Wen, Kimberly M. Christian, P.-Y. Shi, Brady J. Maher, H. Wu, P. Jin, H. Tang, H. Song, and G.-l. Ming, Brain-Region-Specific Organoids Using Mini-bioreactors for Modeling ZIKV Exposure. Cell, 2016. 165(5): p. 1238–1254.

7. Sakaguchi, H., T. Kadoshima, M. Soen, N. Narii, Y. Ishida, M. Ohgushi, J. Takahashi, M. Eiraku, and Y. Sasai, Generation of functional hippocampal neurons from self-organizing human embryonic stem cell-derived dorsomedial telencephalic tissue. Nature Communications, 2015. 6(1): p. 8896.

8. Birey, F., J. Andersen, C.D. Makinson, S. Islam, W. Wei, N. Huber, H.C. Fan, K.R.C. Metzler, G. Panagiotakos, N. Thom, N.A. O’Rourke, L.M. Steinmetz, J.A. Bernstein, J. Hallmayer, J.R. Huguenard, and S.P. Pasca, Assembly of functionally integrated human forebrain spheroids. Nature, 2017. 545(7652): p. 54–59.

9. Xiang, Y., Y. Tanaka, B. Patterson, Y.J. Kang, G. Govindaiah, N. Roselaar, B. Cakir, K.Y. Kim, A.P. Lombroso, S.M. Hwang, M. Zhong, E.G. Stanley, A.G. Elefanty, J.R. Naegele, S.H. Lee, S.M. Weissman, and I.H. Park, Fusion of Regionally Specified hPSC-Derived Organoids Models Human Brain Development and Interneuron Migration. Cell Stem Cell, 2017. 21(3): p. 383–398.e7.

10. Bagley, J.A., D. Reumann, S. Bian, J. Lévi-Strauss, and J.A. Knoblich, Fused cerebral organoids model interactions between brain regions. Nature methods, 2017. 14(7): p. 743–751.

11. Xiang, Y., Y. Tanaka, B. Cakir, B. Patterson, K.Y. Kim, P. Sun, Y.J. Kang, M. Zhong, X. Liu, P. Patra, S.H. Lee, S.M. Weissman, and I.H. Park, hESC-Derived Thalamic Organoids Form Reciprocal Projections When Fused with Cortical Organoids. Cell Stem Cell, 2019. 24(3): p. 487–497.e7.

12. Ogawa, J., G.M. Pao, M.N. Shokhirev, and I.M. Verma, Glioblastoma Model Using Human Cerebral Organoids. Cell Reports, 2018. 23(4): p. 1220–1229.

13. Yoon, S.J., L.S. Elahi, A.M. Pasca, R.M. Marton, A. Gordon, O. Revah, Y. Miura, E.M. Walczak, G.M. Holdgate, H.C. Fan, J.R. Huguenard, D.H. Geschwind, and S.P. Pasca, Reliability of human cortical organoid generation. Nature Methods, 2019. 16(1): p. 75–78.

14. Brandenberg, N., S. Hoehnel, F. Kuttler, K. Homicsko, C. Ceroni, T. Ringel, N. Gjorevski, G. Schwank, G. Coukos, G. Turcatti, and M.P. Lutolf, High-throughput automated organoid culture via stem-cell aggregation in microcavity arrays. Nat Biomed Eng, 2020.

15. Liu, H., Y. Wang, H. Wang, M. Zhao, T. Tao, X. Zhang, and J. Qin, A Droplet Microfluidic System to Fabricate Hybrid Capsules Enabling Stem Cell Organoid Engineering. Advanced Science, 2020. 7(11): p. 1903739.

16. Chen, B., Y. Wu, Z. Ao, H. Cai, A. Nunez, Y. Liu, J. Foley, K. Nephew, X. Lu, and F. Guo, High-throughput acoustofluidic fabrication of tumor spheroids. Lab on a Chip, 2019. 19(10): p. 1755–1763.

17. Guo, F., P. Li, J.B. French, Z. Mao, H. Zhao, S. Li, N. Nama, J.R. Fick, S.J. Benkovic, and T.J. Huang, Controlling cell–cell interactions using surface acoustic waves. Proceedings of the National Academy of Sciences, 2015. 112(1): p. 43–48.

18. Guo, F., Z. Mao, Y. Chen, Z. Xie, J.P. Lata, P. Li, L. Ren, J. Liu, J. Yang, M. Dao, S. Suresh, and T.J. Huang, Three-dimensional manipulation of single cells using surface acoustic waves. Proceedings of the National Academy of Sciences, 2016. 113(6): p. 1522–1527.

19. Ayano, G., Dopamine: receptors, functions, synthesis, pathways, locations and mental disorders: review of literatures. J Ment Disord Treat, 2016. 2(120): p. 2.

20. Nestler, E.J., Is there a common molecular pathway for addiction? Nature neuroscience, 2005. 8(11): p. 1445–1449.

21. Verney, C., Distribution of the catecholaminergic neurons in the central nervous system of human embryos and fetuses. Microscopy Research and Technique, 1999. 46(1): p. 24–47.

22. Qian, X., F. Jacob, M.M. Song, H.N. Nguyen, H. Song, and G.L. Ming, Generation of human brain region-specific organoids using a miniaturized spinning bioreactor. Nat Protoc, 2018. 13(3): p. 565–580.

23. Xiang, Y., Y. Tanaka, B. Cakir, B. Patterson, K.-Y. Kim, P. Sun, Y.-J. Kang, M. Zhong, X. Liu, P. Patra, S.-H. Lee, S.M. Weissman, and I.-H. Park, hESC-Derived Thalamic Organoids Form Reciprocal Projections When Fused with Cortical Organoids. Cell Stem Cell, 2019. 24(3): p. 487–497.e7.

24. Brignani, S. and R.J. Pasterkamp, Neuronal Subset-Specific Migration and Axonal Wiring Mechanisms in the Developing Midbrain Dopamine System. Frontiers in Neuroanatomy, 2017. 11(55).

25. Van den Heuvel, D.M. and R.J. Pasterkamp, Getting connected in the dopamine system. Prog Neurobiol, 2008. 85(1): p. 75–93.

26. Freundlieb, N., C. François, D. Tandé, W.H. Oertel, E.C. Hirsch, and G.U. Höglinger, Dopaminergic Substantia Nigra Neurons Project Topographically Organized to the Subventricular Zone and Stimulate Precursor Cell Proliferation in Aged Primates. The Journal of Neuroscience, 2006. 26(8): p. 2321–2325.

27. Lao, C.L., C.S. Lu, and J.C. Chen, Dopamine D3 receptor activation promotes neural stem/progenitor cell proliferation through AKT and ERK1/2 pathways and expands type-B and-C cells in adult subventricular zone. Glia, 2013. 61(4): p. 475–489.

28. Qian, X., H. Song, and G.L. Ming, Brain organoids: advances, applications and challenges. Development, 2019. 146(8).

29. Zhong, S., S. Zhang, X. Fan, Q. Wu, L. Yan, J. Dong, H. Zhang, L. Li, L. Sun, N. Pan, X. Xu, F. Tang, J. Zhang, J. Qiao, and X. Wang, A single-cell RNA-seq survey of the developmental landscape of the human prefrontal cortex. Nature, 2018. 555(7697): p. 524–528.

30. Lohani, S., A.K. Martig, K. Deisseroth, I.B. Witten, and B. Moghaddam, Dopamine Modulation of Prefrontal Cortex Activity Is Manifold and Operates at Multiple Temporal and Spatial Scales. Cell Rep, 2019. 27(1): p. 99–114.e6.

31. Seong, H.J. and A.G. Carter, D1 Receptor Modulation of Action Potential Firing in a Subpopulation of Layer 5 Pyramidal Neurons in the Prefrontal Cortex. The Journal of Neuroscience, 2012. 32(31): p. 10516–10521.

