## Supporting information for "Controllable Fusion of Human Brain Organoids Using Acoustofluidics"

#### Experimental Section


- **Device fabrication and operation**
- **Simulation of acoustic fields**
- **Fabrication of brain-region specific organoids**
- **Immunofluorescence staining**
- **$Ca^{2+}$ imaging and data processing**
- **Statistical analysis**


#### Supplementary figures


- **Figure S1. Fabrication and characterization of human forebrain organoid**
- **Figure S2. Fabrication and characterization of human midbrain organoid**
- **Figure S3. Characterization of intrinsic heterogeneity of human forebrain organoid**
- **Figure S4. Device image and numerical simulation**
- **Figure S5. Definition of toward and away assembloid.**


#### Supplementary Tables


- **Table S1 Medium composition for human forebrain organoid fabrication**
- **Table S2 Medium composition for human midbrain organoid fabrication**
- **Table S3 Antibody used in immunofluorescence staining**


#### References

#### Experimental Section

**Device fabrication and operation.** The acoustic assembly device consists of six PZTs) embedded into a laser-cut substrate and a cell culture dish. A 9 mm thick acrylic sheet was laser cut into the substrate of the device with an inner chamber of 40mm x 40mm and four small outer chambers for four embedded PZTs. The PZTs (20 x 10 x 3mm, 1MHz resonant frequency) were affixed to the outer chambers with epoxy, and a 3 mm thick acrylic sheet was glued to the substrate bottom to allow the chamber to contain DI water. The opposite two PZTs were wired together to a pair, and two pairs of PZTs were driven independently by two unsynchronized 1MHz RF signals. The RF signals were generated by a function generator (TGP3152, Aim TTi) and amplified by power amplifiers (LZY-22+, Minicircuit) to drive the acoustic assembly device. A cell culture dish (35mm, Greiner Bio) was employed to contain cell solutions and avoid contamination during the acoustic assembly process, the water-filled acrylic chamber was used to guide acoustic wave into the petri dish. In the acoustic assembly experiment, brain region-specific organoids were suspended in phosphate-buffered saline (PBS) supplied with 5% Gel-MA (Sigma-Aldrich) and 1% Irgacure D-2959 (Sigma-Aldrich) were introduced into the acoustic pattern chamber. RF signals (1MHz, 17Vpp) were applied to the PZTs to generate acoustic trapping patterns. After acoustic patterning to ensure the correct positioning of the organoids, the solution was crosslinked for 30 seconds using ultraviolet light (365 nm, 6 mW $cm^{-2}$). The crosslinked solution containing assembloids was transferred to a glass-bottom 24-well plate (MatTek Corporation) for confocal imaging and cultured in the corresponding culture medium.

**Simulation of acoustic fields.** The numerical simulation of the acoustic field in our acoustofluidic manipulation device was conducted using COMSOL Multiphysics 5.2a. To save the computational cost, we simplified the 3D realistic problem to a 2D computational problem. In **Figure S4b**, our numerical model considered the interaction of propagating acoustic waves in the hexagon fluidic chamber, and the three pairs of PZTs were considered as plane acoustic waves radiation boundaries located at the six edges of the hexagon chamber. The 2D model with predefined plane waves radiation boundaries conditions was solved using the frequency domain of predefined pressure acoustics solver in the COMSOL. The corresponding simulation of rotation and 2D manipulation was conducted via modification of amplitudes and phase shift as described in the main text (**Figure S4c**).

**Fabrication of brain-region specific organoids.** Human embryonic stem cell line WA01 was purchased from WiCell. Cells were maintained on Matrigel (Corning) coated 6 well plates and cultured in mTeSR plus medium (Stemcell Technologies) with a medium change every other day in a 37°C, 5% $CO_2$ incubator. WA01 cells were passaged once every week using ReLeSR (Stemcell Technologies). Brain region-specific organoids were fabricated using an adapted protocol.[1] Briefly, 9,000 WA01 cells were suspended in each well of a spheroid formation plate (Corning) to form embryonic bodies (EBs). EBs were then exposed to the forebrain, midbrain brain organoid development strategy respectively following medium composition. The detailed medium composition can be found in Table S1 and S2. Forebrain organoids were embedded in 33% Matrigel (Corning) on day 7, with Matrigel removal on day 14. Midbrain organoids were cultured in suspension and subject to orbital shaker rotation on Day 14. To rotate organoids in an orbital shaker, 6-10 organoids were set in 1 well of a 6-well plate on a standard orbital shaker (Thermo Fisher) and the shaker was set at a speed of 80 rpm.

**Fluorescence protein labeling of midbrain neuron projections:** To visualize midbrain mDA neuron projection from hMO to hFO, 1-week-old hMOs were incubated with a titer > $1X10^9$ vg/mL AAV2-hSyn-mCherry for 24 hours to label hMO neurons. pAAV-hSyn-mCherry was a gift from Karl Deisseroth (Addgene plasmid # 114472; http://n2t.net/addgene:114472; RRID: Addgene_114472). hMOs were allowed 1 week or longer to express mCherry before assembled with hFO to study neuron projection.

**Immunofluorescence staining.** Brain region-specific organoids and assembloids were fixed in 4% paraformaldehyde overnight. The fixed organoids were then washed twice in 1X phosphate-buffered saline (1XPBS) (Gibco) and transferred to 30% sucrose (w/v) to cryoprotect at 4°C overnight. After cryoprotection, the organoids were then embedded in O.C.T. compound (Fisher healthcare) and froze at -80°C. The frozen blocks were then be sectioned into 20μm thickness slices and adhered to charged glass slides (Fisher Scientific). The slices were then blocked with a blocking buffer made with 5% normal goat serum (Abcam) and 0.3 % Triton™ X-100 (Sigma) in 1XPBS (Gibco) for 1 hour at room temperature, followed by primary antibody incubation at 4°C overnight. Primary antibody information can be found in Table S3. The slides were then washed with 1XPBS 3 times and incubated with corresponding secondary antibody at a dilution of 1:500 for 1 hour. The stained slides were washed with 1XPBS 3 times and counterstained with 4',6-diamidino-2- phenylindole (DAPI). To stain proliferative cells with EdU, organoids were labeled with EdU-Alexa 647cell proliferation kit (Invitrogen) according to the manufacturer's protocol. Briefly, forebrain organoids or assembloids were incubated with 10 μM of EdU for 12 hours. The EdU labeled organoids were then fixed in 4% paraformaldehyde overnight and cryoprotected 30% sucrose (w/v) overnight. Organoids were then embedded in O.C.T. compound (Fisher healthcare) and cryo-sectioned into 20μm thickness slices. Sectioned slices were first subjected Click-iT reaction cocktail to label the EdU+ cells with Alexa 647. The labeled slices were then subject to primary and secondary antibody staining as described above.

**$Ca^{2+}$ imaging and data processing.** To image hFO excitatory neuron firing, hFO-hMO assembloids were transfected with CAMKII promoter-driven GCaMP6s calcium sensor expression. AAV.CamKII.GCaMP6s.WPRE.SV40 was a gift from James M. Wilson (Addgene viral prep # 107790-AAV9; http://n2t.net/addgene:107790 ; RRID:Addgene_107790). After transfection at a titer > $1X10^9$ vg/mL for 24 hours, the assembloid were allowed 3 days to fully express the GCaMP6s reporter. The assembloid was then imaged under an Olympus OSR spinning disk microscope at a frame rate of 10 Hz for 5 minutes. The data processing was conducted using a custom processing package based on Python. The processing pipeline included: (i) Motion correction, (ii) source extraction, (iii) activity deconvolution, and (iv) following analysis. The first three steps were performed using CaImAn[2] software to get the deconvoluted $Ca^{2+}$ activity traces. We run CaImAn with default parameters except for some parameters adjusted to fit our records: imaging frame rate in frames per second, half-size of the patches in pixels, overlap between patches in pixels, the number of components per patch, expected half-size of neurons in pixels. The following analysis was performed with extracted $Ca^{2+}$ activity traces. The ISI-distance matrixes were calculated using Pyspike[3] software package.

**Statistical analysis.** The statistics to compare the difference between the two groups were conducted using the students' t-test. Statistical significance was denoted as following: *$p<0.05$, **$p<0.01$, ***$p<0.005$. ****$p<0.001$.

#### Supplementary figures

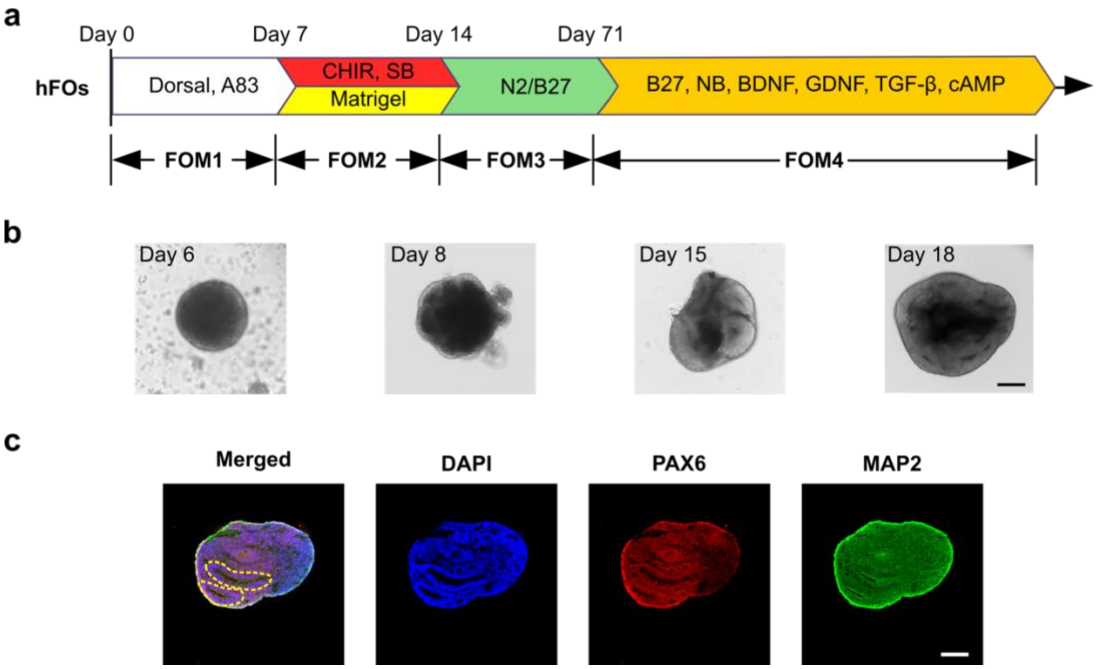


**Figure S1. Fabrication and characterization of human forebrain organoids.** (a) Schematics showing human forebrain organoid fabrication protocol. (b) Human forebrain organoid maturation before the assembly process. (c) Immunofluorescence staining showing human forebrain organoid formation with PAX6+ neural progenitor cells and MAP2+ mature neurons. Scale bar: 200 μm

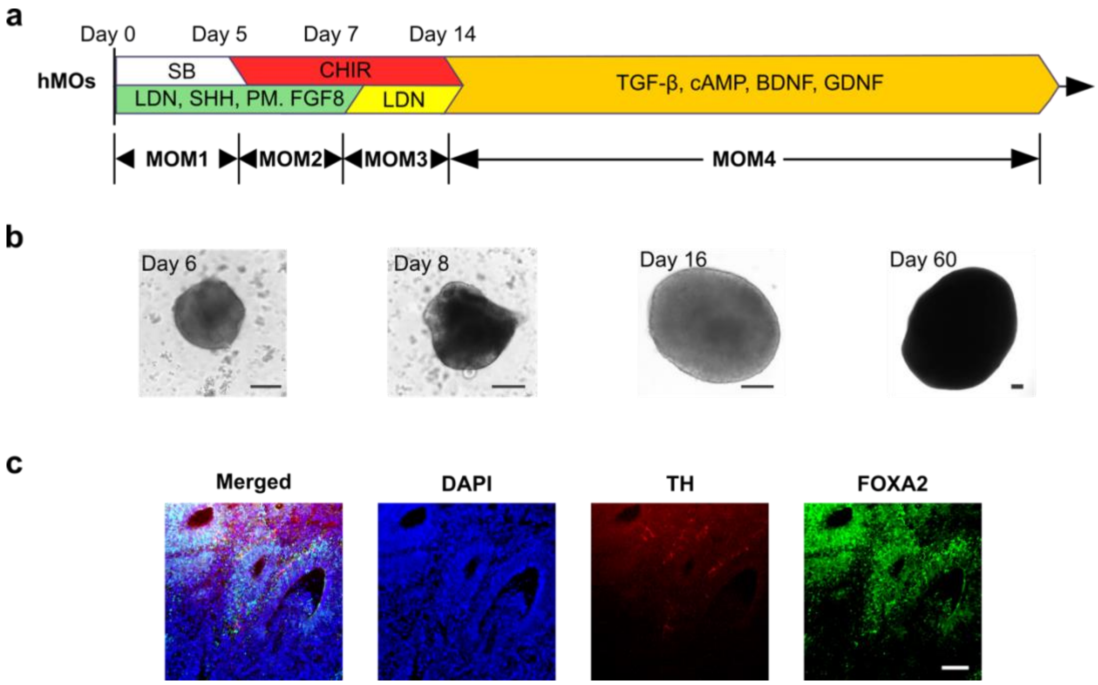


**Figure S2. Fabrication and characterization of human midbrain organoids.** (a) Schematics showing human midbrain organoid fabrication protocol. (b) Human midbrain organoid maturation. Scale bar: 200 μm (c) Immunofluorescence staining showing human midbrain organoid formation with FOXA2+ floor plate progenitor cells and TH+ dopaminergic neurons. Scale bar: 100 μm.

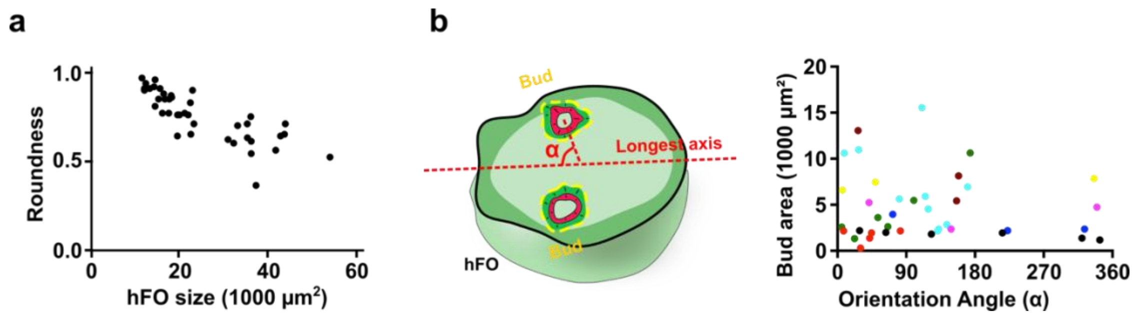


**Figure S3. Characterization of intrinsic heterogeneity of human forebrain organoid.** (a) Size and roundness of human forebrain organoids (n=40). (b) Representative bud localization in forebrain organoids, indicated by size and orientation angle from the longest axis- α. Localization of each bud is indicated by the deviation angle α, which is the orientation angle between the longest axis of the organoid and the line linking the geometric center of the bud and the geometric center of the forebrain organoid. (n=8)

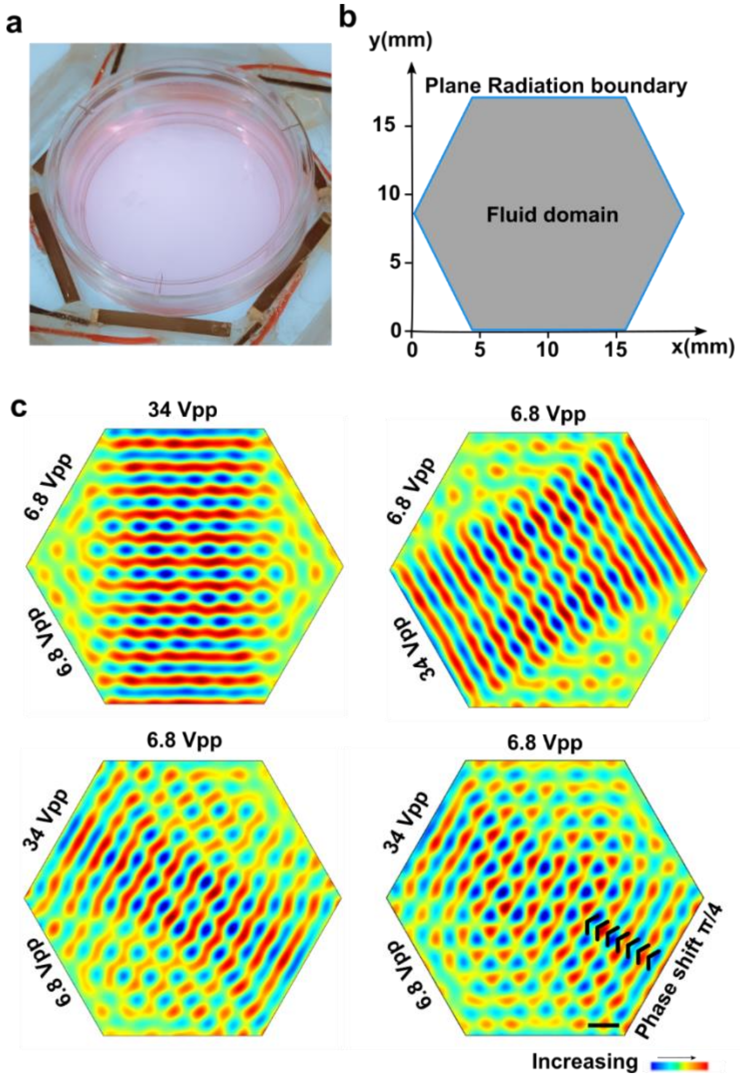


**Figure S4. Device image and numerical simulation** (a) A photo of the acoustofluidic manipulation device. (b) Simulation model consisting of a hexagon fluidic domain and surrounding plane wave radiation boundary accounting for acoustic waves generated by PZTs. (c) A large view of simulation results for acoustofluidic rotational and 2D manipulation. Corresponding to **Figure 2a.** Scale bar: 2 mm

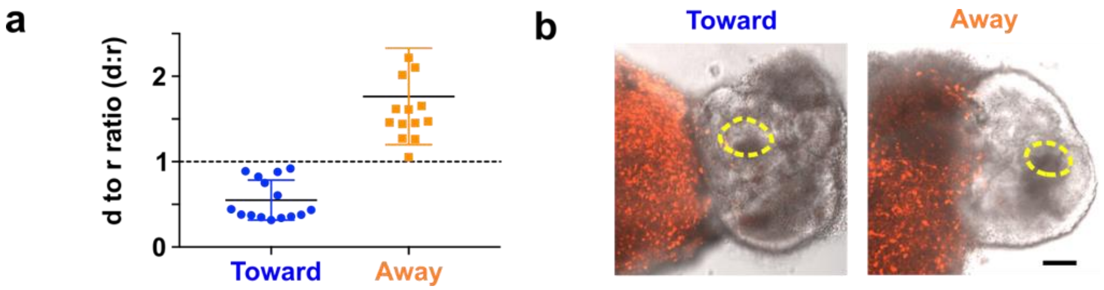


**Figure S5. Definition of toward and away assembloids.** (a) Comparison of hFO bud distance from hMO in the toward and away group. (n=15, p<0.001). (b) Representative images of assembloid in the toward and away group. Buds are marked with yellow dotted circles. Scale bar: 100 μm.

### Supplementary Tables

#### Table S1 Medium composition for human forebrain organoid fabrication

| **Ingredients** | **Concentrations** | **Vendor** | **Catalog#** |
|---|---|---|---|
| Forebrain Medium I (FOM1) | | | |
| DMEM/F12 | 1X | Invitrogen | 11330-032 |
| KOSR | 20% | Invitrogen | 10828028 |
| GlutaMax | 1X | Invitrogen | 35050061 |
| MEM-NEAA | 1X | Sigma | M7145 |
| β-mercaptoethanol | 1X | Sigma | M6250 |
| Dorsomorphin | 2 μM | Stemcell Technologies | 72102 |
| A-83-01 | 2 μM | Stemcell Technologies | 72022 |
| Y-27632 | 5 μM | SelleckChem | S1049 |
| Penn/Strep | 1X | Gibco | 15140122 |
| Forebrain Medium II (FOM2) | | | |
| DMEM/F12 | 1X | Invitrogen | 11330-032 |
| N2 supplement | 1X | Invitrogen | 17502048 |
| GlutaMAX | 1X | Invitrogen | 35050061 |
| MEM-NEAA | 1X | Sigma | M7145 |
| CHIR-99021 | 1 μM | Stemcell Technologies | 72052 |
| SB431542 | 1 μM | Stemcell Technologies | 72232 |
| Penn/Strep | 1X | Gibco | 15140122 |
| Forebrain Medium III (FOM3) | | | |
| DMEM/F12 | 1X | Invitrogen | 11330-032 |
| N2 supplement | 1X | Invitrogen | 17502048 |
| GlutaMAX | 1X | Invitrogen | 35050061 |
| MEM-NEAA | 1X | Sigma | M7145 |
| B27 | 1X | Invitrogen | 17504044 |
| β-mercaptoethanol | 1X | Sigma | M6250 |
| Penn/Strep | 1X | Gibco | 15140122 |
| Insulin | 2.5 μg/mL | Sigma | I9278-5ML |
| Forebrain Medium IV (FOM4) | | | |
| Neuralbasal medium | 1X | Invitrogen | 21103049 |
| GlutaMAX | 1X | Invitrogen | 35050061 |
| MEM-NEAA | 1X | Sigma | M7145 |
| B27 | 1X | Invitrogen | 17504044 |
| Penn/Strep | 1X | Gibco | 15140122 |
| Ascorbic Acid | 0.2 mM | Sigma | 1043003 |
| cAMP | 0.5 mM | Sigma | A9501 |
| BDNF | 20 ng/mL | Peprotech | 450-02 |
| GDNF | 20 ng/mL | Peprotech | 450-10 |

**Table S2 Medium composition for human midbrain organoid fabrication**

| Midbrain Medium I (MOM1) | | | |
|---|---|---|---|
| Ingredients | Concentrations | Vendor | Catalog# |
| DMEM/F12 | 1X | Invitrogen | 11330-032 |
| KOSR | 20% | Invitrogen | 10828028 |
| GlutaMax | 1X | Invitrogen | 35050061 |
| MEM-NEAA | 1X | Sigma | M7145 |
| β-mercaptoethanol | 1X | Sigma | M6250 |
| LDN-193189 | 100 nM | Stemcell Technologies | 72147 |
| SB-431542 | 10 μM | Stemcell Technologies | 72232 |
| SHH | 100 ng/mL | Peprotech | 100-45 |
| Purmorphamine | 2 μM | Stemcell Technologies | 72202 |
| FGF8 | 100 ng/mL | Peprotech | 100-25A |
| Penn/Strep | 1X | Gibco | 15140122 |
| Midbrain Medium II (MOM2) | | | |
| DMEM/F12 | 1X | Invitrogen | 11330-032 |
| N2 supplement | 1X | Invitrogen | 17502048 |
| GlutaMAX | 1X | Invitrogen | 35050061 |
| LDN-193189 | 100 nM | Stemcell Technologies | 72147 |
| CHIR-99021 | 3 μM | Stemcell Technologies | 72052 |
| SHH | 100 ng/mL | Peprotech | 100-45 |
| Purmorphamine | 2 μM | Stemcell Technologies | 72202 |
| FGF8 | 100 ng/mL | Peprotech | 100-25A |
| Penn/Strep | 1X | Gibco | 15140122 |
| Midbrain Medium III (MOM3) | | | |
| DMEM/F12 | 1X | Invitrogen | 11330-032 |
| N2 supplement | 1X | Invitrogen | 17502048 |
| GlutaMAX | 1X | Invitrogen | 35050061 |
| MEM-NEAA | 1X | Sigma | M7145 |
| LDN-193189 | 100 nM | Stemcell Technologies | 72147 |
| CHIR-99021 | 3 μM | Stemcell Technologies | 72052 |
| Penn/Strep | 1X | Gibco | 15140122 |
| Midbrain Medium IV (MOM4) | | | |
| Neuralbasal medium | 1X | Invitrogen | 21103049 |
| GlutaMAX | 1X | Invitrogen | 35050061 |
| MEM-NEAA | 1X | Sigma | M7145 |
| B27 | 1X | Invitrogen | 17504044 |
| Penn/Strep | 1X | Gibco | 15140122 |
| Ascorbic Acid | 0.2 mM | Sigma | 1043003 |
| cAMP | 0.5 mM | Sigma | A9501 |
| BDNF | 20 ng/mL | Peprotech | 450-02 |
| GDNF | 20 ng/mL | Peprotech | 450-10 |

**Table S3 Antibody used in immunofluorescence staining**

| Antigen | Host | Vendor | Catalog# | Dilution |
|---|---|---|---|---|
| PAX6 | Rabbit | Biolegend | 901301 | 1:500 |
| MAP2 | Chicken | Millipore | AB5543 | 1:500 |
| TH | Rabbit | Millipore | AB152 | 1:200 |
| FOXA2 | Goat | R&D | AF2400 | 1:200 |

#### References


[1] X. Qian, F. Jacob, M. M. Song, H. N. Nguyen, H. Song, G.-l. Ming, Nature Protocols 2018, 13, 565.

[2] A. Giovannucci, J. Friedrich, P. Gunn, J. Kalfon, B. L. Brown, S. A. Koay, J. Taxidis, F. Najafi, J. L. Gauthier, P. Zhou, B. S. Khakh, D. W. Tank, D. B. Chklovskii, E. A. Pnevmatikakis, Elife 2019, 8.

[3] T. Kreuz, D. Chicharro, C. Houghton, R. G. Andrzejak, F. Mormann, J Neurophysiol 2013, 109, 1457; T. Kreuz, M. Mulansky, N. Bozanic, J Neurophysiol 2015, 113, 3432.
